# Hungarian Grey cattle grazing outperforms mowing during wet meadow restoration following plantation forest clear-cut

**DOI:** 10.1101/2023.04.28.538658

**Authors:** Katalin Szitár, Melinda Kabai, Zita Zimmermann, Gábor Szabó, Bruna Paolinelli Reis, László Somay

**Affiliations:** ’Lendület’ Landscape and Conservation Ecology, Institute of Ecology and Botany, Centre for Ecological Research, Vácrátót, Hungary; Restoration Ecology, Institute of Ecology and Botany, Centre for Ecological Research, Vácrátót, Hungary; Doctoral School of Environmental Sciences, Hungarian University of Agriculture and Life Sciences, Gödöllő, Hungary; cE3c, Centre for Ecology, Evolution and Environmental Changes & CHANGE - Global Change and Sustainability Institute, Faculdade de Ciências, Universidade de Lisboa, Campo Grande, Lisboa, Portugal; ’Lendület’ Ecosystem Services, Institute of Ecology and Botany, Centre for Ecological Research, Vácrátót, Hungary

**Keywords:** functional trait, Pannonian ecoregion, temperate wet grassland, traditional cattle breed, Natura 2000 site

## Abstract

Land-use change and ecological invasion are two main drivers of biodiversity loss, and the restoration of semi-natural wet grasslands is needed to tackle invasive species and re-establish grassland biodiversity on former forest plantations. This study tested the effectiveness of two widely used management techniques (grazing by traditional Hungarian Grey cattle and mowing once a year in August) as a restoration method of wet meadows in a former forest plantation invaded by goldenrod species in Central Hungary. We compared the vegetation composition of grazed, mowed, and reference areas with semi-natural wet meadow vegetation based on plant biomass, species richness and cover of species groups of species origin, life span, growth form, and social behaviour types of Borhidi determining the grazing value and the nature conservation value of the grasslands. We found that grazing by Hungarian Grey cattle resulted in a vegetation that was more similar to the reference wet meadows than mowing once a year in late summer. Grazing was superior to mowing in terms of goldenrod control, total species richness and cover, as well as the abundance of natives, perennials, herbs, and legumes. However, in the grazed area, we detected more disturbance-tolerant and annual species than in the mowed area. Despite the improved vegetation condition in the grazed area, we identified substantial disparities between the grazed and reference areas after three years of grazing. Based on our results, we advise using continuous extensive grazing to restore and maintain semi-natural wet meadows.

## 1. Introduction

Land-use change and ecological invasion are among the main drivers of biodiversity loss and habitat degradation (IPBES, 2019). In the last centuries, afforestation of natural and semi-natural grassland ecosystems has been a common practice worldwide (Koyama and Uchida, 2022). Palearctic grasslands support high biodiversity with a high proportion of grassland specific and endemic flora (Dengler et al., 2014). They also provide ecosystem services such as livestock production, and pollination services that plantation forests cannot do (Naidoo et al., 2008, Martínez et al., 2009).

Forest plantations are typically created with few or single tree species covering large areas and therefore, have serious landscape-scale homogenizing effects (Hu et al., 2022). The conversion of natural grasslands represents major abiotic changes from herbaceous to arboreal vegetation (Szitár et al., 2014), altering hydrologic, light, and nutrient conditions (Buytaert et al., 2007). Soil preparation before tree planting is also a major driver to loss of grassland-specific flora (Rédei et al., 2020), and their ecosystem functions and services (Sandoval López et al., 2020). Because of soil disturbance and missing competing herb layer, tree plantations can be hot-spots of plant invasion (Csecserits et al., 2016). In Central Europe, alien goldenrod species (*Solidago gigantea* and *S. canadensis*) have been invaded large areas in both semi-natural and transformed ecosystems (Szymura et al., 2016). Goldenrod invasion has been shown to result in decrease in plant, wild pollinator (Fenesi et al., 2015), bird species richness (Skórka et al., 2010), an alteration of primary productivity (Vanderhoeven et al., 2006), and spontaneous succession (Gusev, 2015). After forest clear-cut, invasive species such as goldenrod species often remain permanently in the vegetation impeding spontaneous regeneration (Szitár et al., 2016).

The restoration of semi-natural grasslands is highly needed to re-establish grassland biodiversity and related ecosystem functions on former forest plantations (Zaloumis and Bond, 2011). In the course of restoration, both abiotic limitations (e.g. microsites for species establishment) and biotic constraints (e.g. limitations of plant propagules, and species dominance) have to be overcome (Bakker and Berendse, 1999). Restored grasslands also need post-restoration management to maintain the favourable ecological status (Török et al., 2021). Mowing and grazing are the most frequently methods used for removing aboveground biomass and to enhance species diversity of grasslands during restoration (Collins et al., 1998). In their meta-analysis, Tälle et al. (2016) found that grazing was more favourable on conservation values than mowing in case of most grassland types. Given the limited financial resources, the most efficient treatments should be used to optimise grassland management.

The preservation of traditional land use practices has been widely advised to meet biodiversity and production goals in sustainable and multifunctional grassland systems (Isselstein et al., 2007). Traditional livestock breeds are more adapted to the local environment with a stronger resistance to diseases as they have lived in the local environment for generations (Roosen et al., 2005). They are also better suited to utilise grasslands under extensive grazing regime (Isselstein et al., 2007). They maintain higher species diversity at the landscape scale and are able to suppress noxious weeds; therefore, they should be prioritised over commercial livestock breeds (Kovácsné Koncz et al., 2020).

In our study, we tested the effectiveness of two widely used management technique (grazing and mowing) on the restoration of wet meadows invaded by goldenrod species in a former forest plantation (*Populus x euramericana*) in Central Hungary. We tested whether mowing once a year or extensive grazing by traditional Hungarian Grey cattle results in the restoration of the vegetation of wet meadows in terms of biomass, species richness and cover of species groups determining the grazing value and the nature conservation value of the grasslands. Our specific hypotheses were the following: (1) Hungarian Grey cattle grazing results in more a favourable vegetation based on forage quality and quantity. (2) Hungarian Grey cattle grazing outperforms the mowing treatment in terms of goldenrod control. (3) Hungarian Grey cattle grazing results in a vegetation more similar to the semi-natural reference area compared to the mowing in terms of species characteristics (origin, life span, growth form, and social behaviour types of Borhidi, 1995).

## 2. Methods

### 2.1. Study site

Our study was conducted in the Zámoly Basin in Central Hungary (Fig. 1). The area is part of the Natura 2000 Special Protection Area (HUDI30002). The basin is situated at the foothills of the Vértes Mountain. The climate is continental with a sub-Mediterranean influence. The mean annual temperature is 9.8-10 °C, and the mean annual precipitation is 550-600 mm (Uj et al., 2012). The soils can be characterised mainly as chernozem and meadow soils (Pásztor et al., 2018), while the dominant semi-natural vegetation types include tussock-forming *Festuca*- and *Stipa*-dominated dry grasslands, marshes, rich fens, *Molinia* meadows, mesotrophic wet meadows, and hay meadows (Pallag, 2009).

**Fig. 1.**
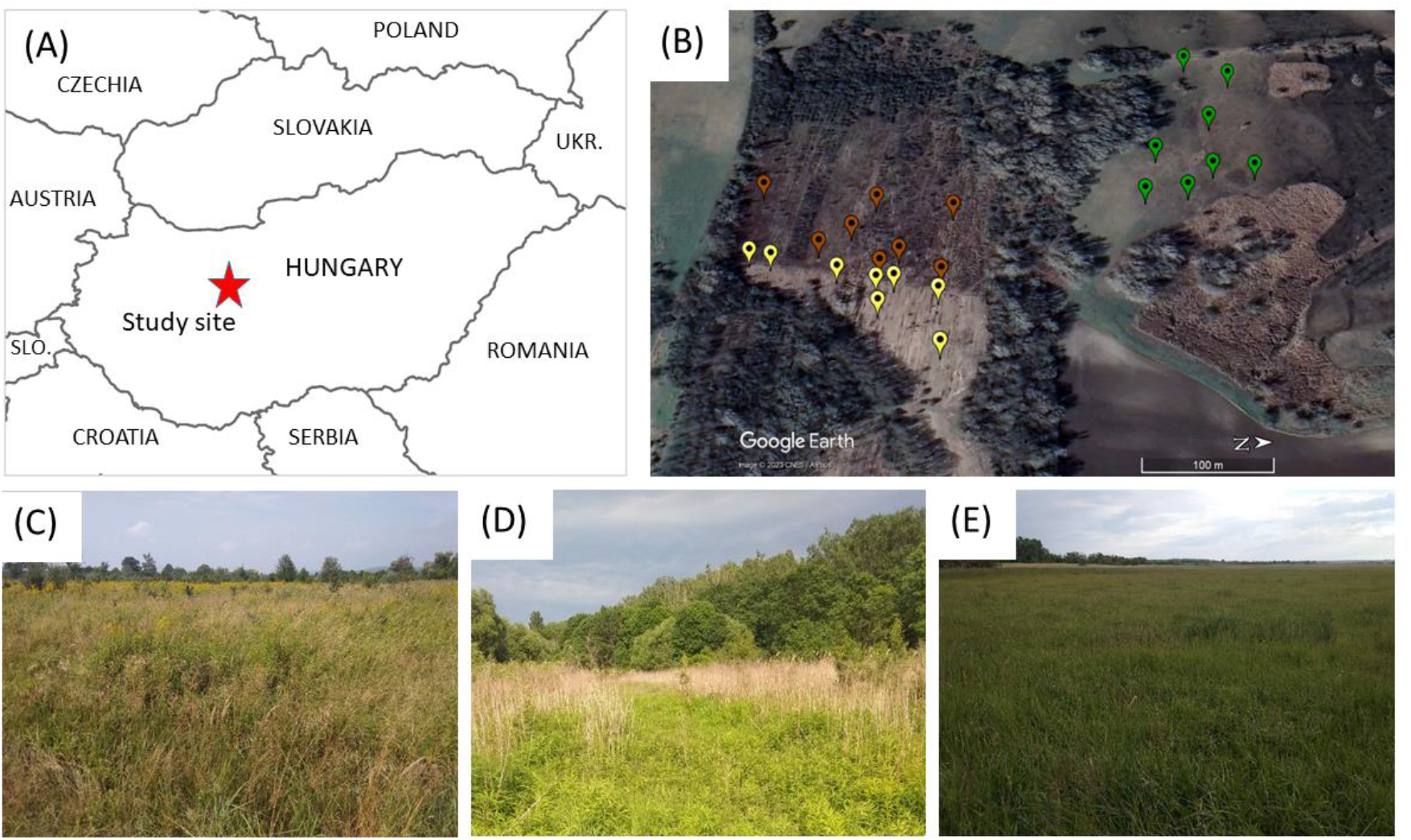
(A) Map of Eastern Europe with the study site (red star), (B) map of the study site with the three vegetation types (brown: Grazed, yellow: Mowed, and green: Reference), photo of the (C) Grazed, (D) Mowed, and (E) Reference sampling sites.

Our study site is located in the Rétföld in the North-eastern part of the Zámoly Basin (47.376° N, 18.487° E). Based on historic maps, the original semi-natural vegetation of the 7-hectares site was wet meadow that was converted into arable land by the end of the 19^th^ century, and later transformed into a non-native poplar plantation (*Populus x euramericana*). The plantation was invaded heavily by giant goldenrod (*Solidago gigantea*). The plantation was cleared from the trees in 2017 to allow the regeneration of wet meadow vegetation. In the same year, the site was divided into two parts with a fence. In a 5-hectares part (hereafter Grazed), grazing by the traditional Hungarian Grey cattle was started. A herd of 700 Hungarian Grey cattle grazes the vegetation with free roaming controlled by a system of electric fences between the second half of July and October each year. The other 2-hectares part of the site (hereafter Mowed) was excluded from grazing, and here, a yearly mowing (in August) is implemented. As a reference (hereafter Reference), we designated an area of wet meadows North to the Grazed site. The vegetation of the Reference is natural wet meadow (*Agrostio-Caricetum distantis*) and was not invaded by giant goldenrod. The Reference is grazed by the same herd of Hungarian Grey cattle as the Grazed site, and is mowed once a year in August.

### 2.2. Vegetation sampling and species functional groups

In the three parts, we sampled the vegetation in June 2020 before grazing and mowing. We made coenological relevés in altogether 24 2 m x 2 m quadrats to characterise the vegetation composition. We estimated the aboveground percentage cover of each vascular plant species.

The species were grouped based several functions reflecting the ecological condition and pasture quality of the vegetation. Species were assigned to functional groups based on their origin (native vs adventive), life span (annual, biennial, or perennial), and growth form (graminoid, legume, herb, or phanerophyte). Based on Borhidi (1995), we also used social behaviour types (hereafter SBT) reflecting the ecological role of species played in the vegetation. We applied the following SBT categories: competitor (dominant species of natural communities), specialist (stress tolerant with narrow stress tolerance and low competitiveness), generalist (stress tolerant with wide stress tolerance), natural pioneers (plants associated with habitats in initial successional stage after natural disturbances), disturbance tolerant (plants of natural habitats with anthropogenic disturbance), weed (native anthropogenic species), ruderal competitor (native community-forming dominant weeds), and alien competitor (habitat transforming invasive species). The number and cover of specialists, natural pioneers, and alien competitors were negligible, therefore we did not analyse them. Group values were collected from the PADAPT database (Sonkoly et al., 2022), the Flora database 1.2. (Horváth et al., 1995), and the identification guide of the Hungarian flora (Király, 2009). For details of species list, trait values and average cover of the species in each management type, see Table S1.

Next to each quadrat, we also made biomass sampling by cutting the vegetation at the height of 1 cm from the soil surface in three 25 cm x 25 cm quadrats on three sides of the coenological quadrats. The biomass samples were sorted as graminoids (*Poaceae* and *Cyperaceae*), herbs (including legumes, trees and shrubs) and goldenrod fractions, and then dried at 65 °C for 48 h. The dry weight of the three fractions was measured with an accuracy of 0.01 g.

### 2.3. Statistical analyses

For each relevé, we calculated the total species richness and the number of species assigned to each functional group category. We also calculated the relative cover of each functional group category by dividing the percentage cover of the species assigned to the category by the total vegetation cover. To disentangle the effect of the management on the abundance of goldenrod and other species, we did not consider goldenrod in the analyses of total richness, total cover, and functional groups but analysed it separately. We tested the effects of management type (Grazed, Mowed, and Reference) on the total species richness, and the richness of functional groups by using general linear models with Poisson error distribution and log link function (lme4 package; Bates et al., 2015). We tested the models for overdispersion with the DHARMa package (Hartig, 2022). We used linear models for biomass and cover data (lme4 package; Bates et al., 2015) with logit transformation of the cover values to fulfil test assumptions (Fox and Weisberg, 2019). The model residuals were visually inspected for heteroscedasticity by checking residuals against fitted values. We derived the adjusted R-squared of the models to estimate the variance explained by the models using the glmtoolbox (Vanegas et al., 2022). In case of significant management type effect, Tukey post hoc pairwise comparisons were made with the multcomp package (Lenth et al., 2018), and were presented in the figures.

All analyses were conducted in the R environment (R Core Team, 2022).

## 3. Results

### 3.1. Species richness and total cover

We recorded altogether 102 vascular plant species in the three sampling sites. The Grazed plots had the highest species richness with 66 species, followed by the Mowed and Reference sampling quadrats with 52 species each. We found that the mean species richness was not significantly different in the Grazed and Reference plots (Table 1, Fig. 2A). In contrast, the Mowed plots harboured 25% less species on average compared to the Reference plots. The average total cover was 133.4 and 134.0 in case of Grazed and Mowed plots, respectively, exceeding the total cover of Reference plots. However, the total cover of plant species excluding goldenrod was the highest in the Reference plots that were free of goldenrod invasion (Fig. 2B). We also found that the cover of goldenrod was 45% lower in the Grazed plots than in the Mowed plots.

**Table 1.** ANOVA table of the linear models and generalized linear models for species richness, cover, and biomass data. Species richness and species cover data were analysed excluding goldenrod. Herb biomass included the biomass of legume and phanerophyte species. Reference data were omitted in the analyses of goldenrod biomass and cover, as they did not contain the species. P values less than 0.05 are shown in bold.

| Response variable | df | F or $\chi^2$ | P | adjusted R<sup>2</sup> |
| --- | --- | --- | --- | --- |
| Species richness | 2 | 12.91 | **0.002** | 0.254 |
| Species cover | 2,21 | 3.61 | **0.045** | 0.185 |
| Goldenrod cover | 1,14 | 8.27 | **0.012** | 0.326 |
| Graminoid biomass | 2,21 | 0.99 | 0.388 | 0.086 |
| Herb biomass | 2,21 | 3.68 | **0.043** | 0.189 |
| Goldenrod biomass | 1,14 | 5.89 | **0.029** | 0.246 |

**Table 2.** ANOVA table of the linear models and generalized linear models. Reference data were omitted in the analyses of biennial and phanerophyte species, as they did not contain any species of these categories. P values less than 0.05 are shown in bold.

| Response variable | Richness |  |  |  | Cover |  |  |  |
| --- | --- | --- | --- | --- | --- | --- | --- | --- |
| | df | $\chi^2$ | P | adj-R<sup>2</sup> | df | F | P | adj-R<sup>2</sup> |
| *Origin* |  |  |  |  |  |  |  |  |
| Native | 2 | 13.27 | **0.001** | 0.248 | 2,21 | 0.72 | 0.497 | 0.065 |
| Exotic | 2 | 3.24 | **0.198** | 0.153 | 2,21 | 0.72 | 0.497 | 0.065 |
| *Life span* |  |  |  |  |  |  |  |  |
| Perennial | 2 | 11.91 | **0.003** | 0.243 | 2,21 | 12.63 | **<0.001** | 0.503 |
| Biennial* | 1 | 0.11 | 0.739 | 0.001 | 1,14 | 0.04 | 0.840 | 0.003 |
| Annual | 2 | 23.86 | **<0.001** | 0.495 | 2,21 | 5.19 | **0.015** | 0.267 |
| *Growth form* |  |  |  |  |  |  |  |  |
| Graminoid | 2 | 3.64 | 0.162 | 0.122 | 2,21 | 0.56 | 0.579 | 0.051 |
| Herb | 2 | 9.82 | **0.007** | 0.230 | 2,21 | 15.14 | **<0.001** | 0.552 |
| Legume | 2 | 7.15 | **0.028** | 0.153 | 2,21 | 2.84 | 0.081 | 0.138 |
| Phanerophyte* | 1 | 1.00 | 0.317 | 0.080 | 1,14 | 0.32 | 0.580 | 0.022 |
| *Social behaviour type* |  |  |  |  |  |  |  |  |
| Competitor | 2 | 9.33 | **0.009** | 0.391 | 2,21 | 0.59 | 0.563 | 0.053 |
| Generalist | 2 | 4.39 | 0.111 | 0.009 | 2,21 | 1.71 | 0.205 | 0.058 |
| Disturbant tolerant | 2 | 10.93 | **0.004** | 0.268 | 2,21 | 1.95 | 0.167 | 0.076 |
| Weed | 2 | 7.07 | **0.029** | 0.368 | 2,21 | 6.55 | **0.006** | 0.325 |
| Ruderal competitor | 2 | 6.71 | **0.035** | 0.185 | 2,21 | 2.43 | 0.113 | 0.110 |

**Fig. 2.**
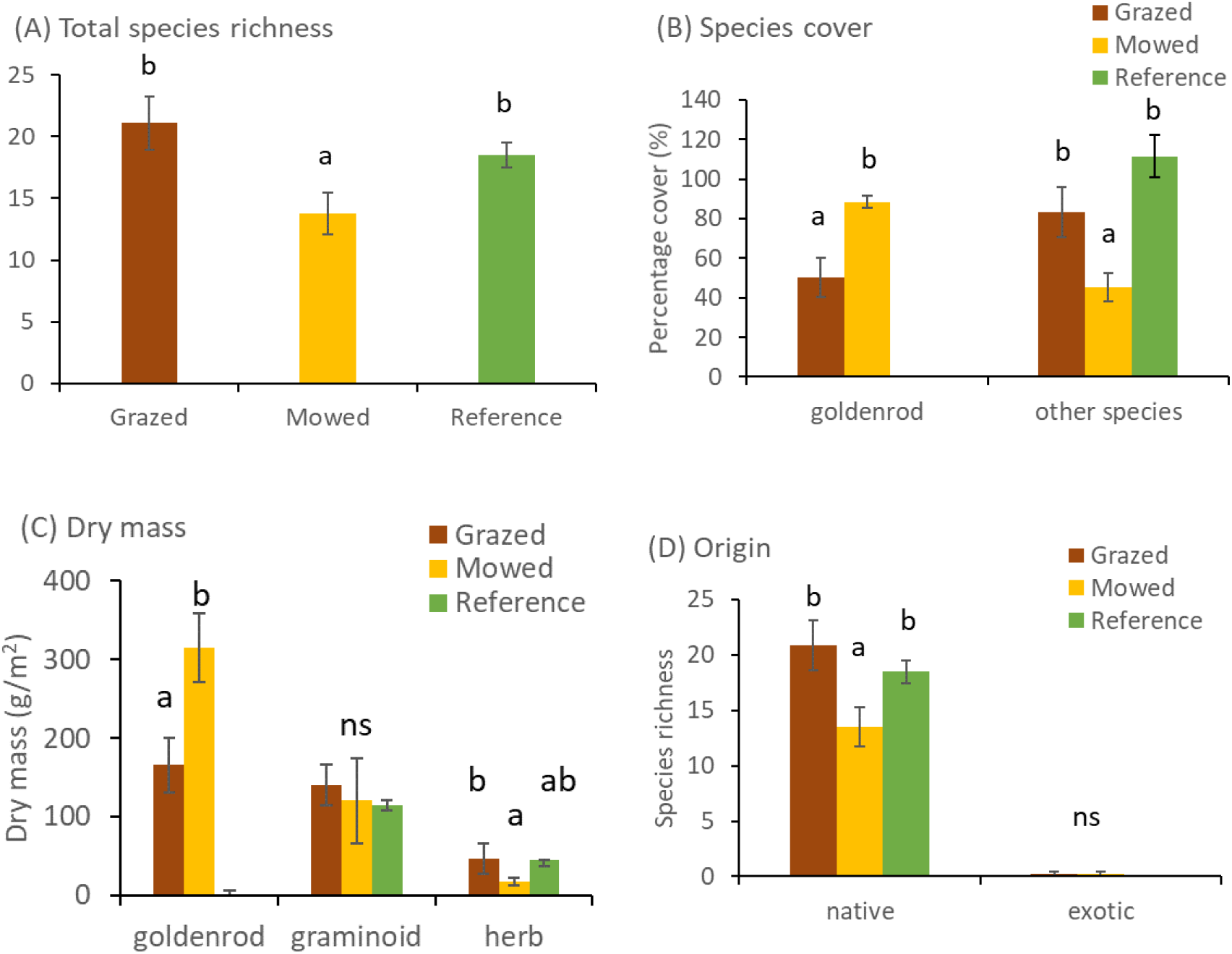
(A) Total species richness (Mean ± SEM), and (B) cover (Mean ± SEM) of goldenrod (*Solidago gigantea*) and other species, (C) mean dry mass of goldenrod, graminoids, and herbs, and (D) the richness of native and exotic species in the three management types. Different letters denote statistical difference of the management types.

### 3.2. Biomass production

The dry mass of goldenrod was 48% lower in the Grazed plots than the Mowed plots (Fig. C). There were no differences among the treatment types in the biomass of graminoid species. Herb species (including legumes and phanerophytes) had higher biomass in the Grazed plots and the lower in the Mowed plots but they did not differ from the Reference.

### 3.3. Origin of species, life span, life form, and social behaviour types

The number and cover of exotic species excluding goldenrod was minimal in every treatment types (Fig. 2D). We found that the number of native species showed a similar pattern to that of the total species richness with a significantly lower richness in the Mowed plots compared to both the Grazed and Reference plots. Perennial species dominated the vegetation with over 10 species and 90% cover in all management types (Fig. 3A and B). Both the richness and the cover of perennial species were the highest in the Reference plots, and the lowest in the Mowed plots, whereas the richness of annual species in the Grazed plots exceeded that of the other treatment types (Fig. 3C and D). Similar to the biomass, the richness and cover of graminoid species and were not significantly different among the treatment types. Herbs were more species rich in the Grazed and Reference plots compared to the Mowed treatment plots but the cover of herbs were equally low in the Grazed and Mowed plots compared to the Reference. We found that the Mowed plots had less legume species than the other two treatment types but this difference was not detected in the case of species cover. We found phanerophyte species in the Grazed and Mowed plots with similar richness and cover but these were missing from the Reference plots. The species richness showed differences in four social behaviour types. We found the highest number of competitor and generalist species in the Reference plots, whereas Grazed plots were the richest in disturbance tolerants, weeds and ruderal competitors (Fig. 3E). The cover of social behaviour types were similar in the three treatments, except for weeds that were almost missing in the Reference plots (Fig. 3F).

**Fig. 3.**
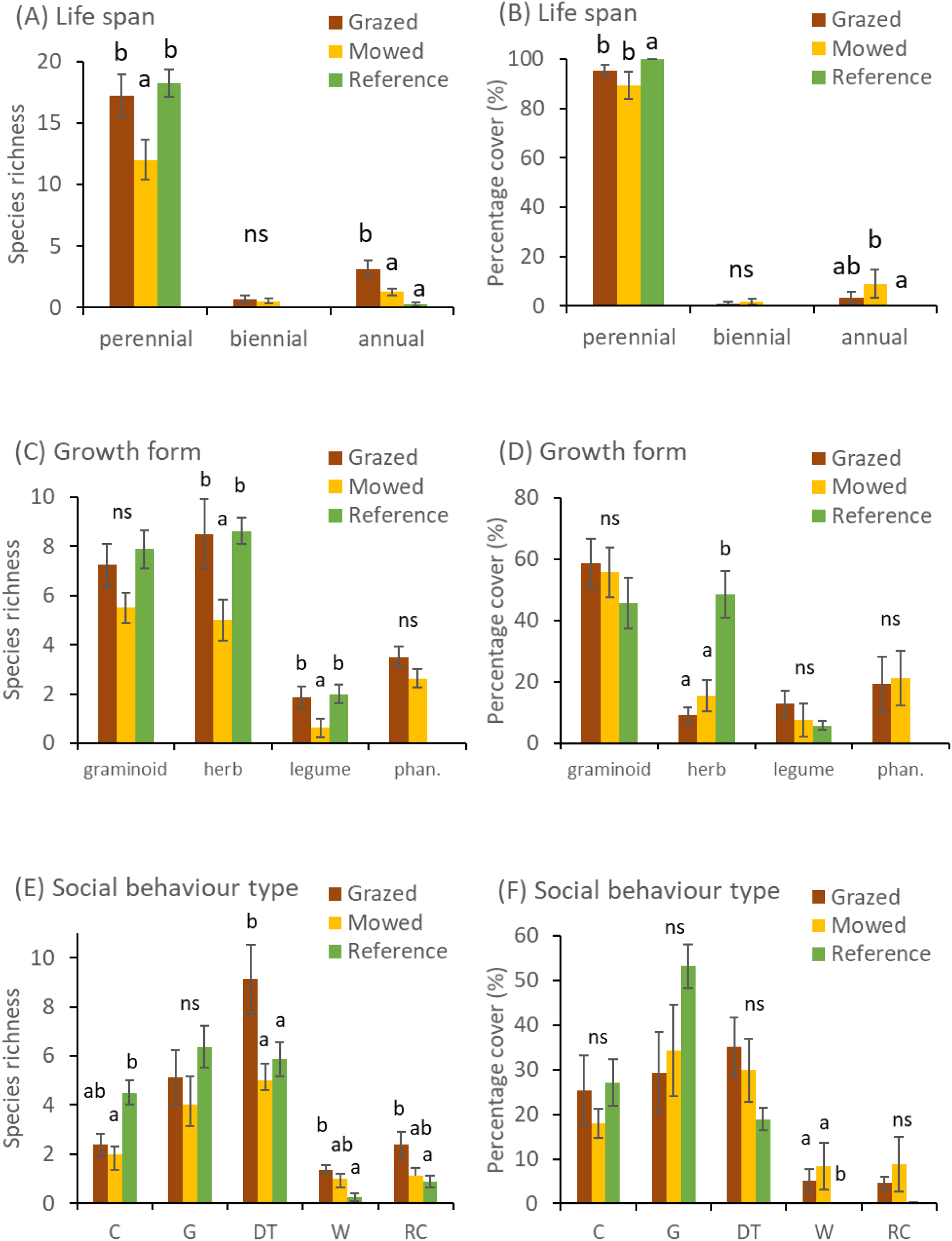
Species richness (Mean ± SEM), and cover (Mean ± SEM) of species with different life span (A and B), growth form categories (C and D), and Social behaviour types (Borhidi, 1995; E and F). Abbreviations: phan: phanerophyte (trees and bushes), C: competitor, G: generalist, DT: disturbance tolerant, W: weed, RC: ruderal competitor. Different letters denote statistical difference of the management types within each category, ns stands for no significant difference among the treatment groups.

## 4. Discussion

We found that grazing by Hungarian Grey cattle resulted in a vegetation that was more similar to the reference wet meadows than mowing once a year in late summer. Grazing was superior to mowing in terms of goldenrod control, total species richness and cover, as well as the abundance of natives, perennials, herbs, and legumes. However, in the grazed area, we detected more disturbance-tolerant and annual species than in the mowed area. Grazing also resulted in higher dry mass of herbs, excluding goldenrod, than mowing. Despite the improved vegetation conditions in the grazed region, we identified substantial disparities between the grazed and reference areas after three years of grazing.

### 4.1. Forage quality and quantity

For sustainability, forage production is of critical importance in grazing systems (Isselstein et al., 2007). If we exclude goldenrod biomass, total biomass of plant species reached in the grazed and mowed areas that of the reference, showing a rapid successful regeneration of forage quantity. This contradicts the findings of multiple research that demonstrate that both grazing and mowing reduce aboveground biomass (Török et al., 2011). We assume that our extensive grazing and mowing contributed to high biomass production, similar to the results of Martin and Wilsey (2006) who showed that grazing intensity had a negative effect on plant biomass. Cattle grazing resulted in more a favourable vegetation based on forage quality than mowing. Although, we did not detect any differences in grass abundance among the three treatments, whereas herb biomass, and the richness of legumes and other herbs were higher in grazed plots than in mowed plots. High legume abundance has been shown to boost the forage’s digestibility and crude protein content (White and Wight, 1984), and many herb species have therapeutic properties, making them useful not just for nature conservation but also for animal health (Mirzaei-Aghsaghali, 2012).

### 4.2. Goldenrod control

We found that extensive grazing with traditional Hungarian Grey cattle resulted in a significant reduction of cover and biomass of giant goldenrod (*S. gigantea*) compared to the mowing treatment. Nagy et al. (2020) found that goldenrod density could be decreased successfully using both grazing and mowing. Earlier studies also showed also stated that grazing with Hungarian Grey cattle was an efficient tool to suppress goldenrod invasion (Malatinszky et al., 2013; Visnyovszky, 2017). According to the biotic resistance hypothesis (Levine, 2000), communities with high species richness at small spatial scales are generally more resistant to plant invasion. In our study, Hungarian Grey cattle grazing supported the highest species diversity showing the potential for a higher resistance to further goldenrod establishment (Nagy et al., 2020).

It is important to note, however, that neither grazing nor mowing resulted in a complete removal of goldenrod populations. Total elimination of goldenrod through mowing and grazing is rarely feasible (Szymura et al., 2022) but they are suitable to reduce goldenrod cover to a low level even within three years (Hall et al., 2022). Subsequent management is also required to prevent goldenrod recovery (Szymura et al., 2022) as the species has outstanding seed dispersal abilities, high seed production and effective vegetative reproduction (Weber, 2011).

### 4.3. Vegetation composition

We discovered that grazing produced vegetation that was more similar to the reference wet meadows than plots with mowing once a year in late summer. Grazing was superior to mowing in terms of the species richness of natives, annuals, perennials, herbs, and legumes. Based on the meta-analysis of Tälle et al. (2016), grazing was also more favourable to the vegetation condition of wet semi-natural grasslands than mowing. Grazing and mowing create different types of disturbance influencing subsequent vegetation changes (Nagy et al., 2020). Mowing was shown to cause vegetation homogenisation when implemented within a short period of time with large machinery (Zechmeister et al., 2003). The higher plant diversity under grazing may be the result of elevated microsite diversity (Tölgyesi et al., 2015), selective grazing of strong competitors (Hartnett et al., 1996), and gap creation by trampling (Loucougaray et al., 2004).

Mowing and grazing did not differ in the effect on the cover of species groups of life span, growth form, and social behaviour types. The absence of these effects may be due to the preferential grazing of dominant species equating species abundances in the vegetation (Hartnett et al., 1996). We found that both annuals and disturbance tolerant species disturbance tolerant species increased in richness in response to grazing similar to Nagy et al. (2020). Kiss et al. (2011) also showed that disturbance tolerants and ruderal competitors accumulate near animal husbandry sites with intensive trampling and grazing.

Despite the improved vegetation conditions in the grazed region, we identified substantial disparities among the grazed, mowed, and reference areas after three years of introduction of grazing and mowing. The cover of perennial and herb cover in the reference area still exceeded that in the grazed and mowed sites. All three vegetation types were dominated by disturbance tolerant and generalist species but reference areas harboured a higher number of competitor species characteristic to wet meadows and less weeds. The length of time since restoration started is an important determinant for vegetation regeneration (Kettenring and Adams, 2011). Although cattle grazing strongly affects vegetation composition even in the short run (Török et al., 2014), the complete regeneration of wet grasslands in terms of vegetation composition may take several decades (Moeslund et al., 2022).

## Conclusions

In this study, we showed that the restoration of wet meadows in the place of clear-cut forest plantation was more successful with extensive grazing with Hungarian Grey cattle than with mowing once a year in August, although, the vegetation regeneration was not complete after three years of management even in the case of grazing treatment. Based on our results, we suggest the use of continuous extensive grazing for the restoration and maintenance of semi-natural wet meadows.

## Acknowledgements

This study was supported by the Hungarian Széchenyi 2020 Programme (VP3-16.1.1-4.1.5 No. 3043361958), and the Hungarian National Research, Development and Innovation Office (NKFIH KKP133839).

