## Supplementary Table S1 for "Hungarian Grey cattle grazing outperforms mowing during wet meadow restoration following plantation forest clear-cut"

Supplementary information

Table S1. of the analysed plant species with their analysed characteristics, their frequency and average cover in the three management types. Species were assigned to functional groups based on their origin (native vs adventive), life span (annual, biennial, or perennial), growth form (graminoid, legume, herb, or phanerophyte), and social behaviour types (SBT). Abbreviations: Ref: Reference, C: competitor, S: specialist, G: generalist, NP: DT: disturbance tolerant, RC: ruderal competitor, AC: alien competitor. NA denotes missing values.

| Species name | Origin | Life span | Growth form | SBT | Frequency | | | Mean cover | | |
|---|---|---|---|---|---|---|---|---|---|---|
| | | | | | Grazed | Mowed | Ref. | Grazed | Mowed | Ref. |
| **Acer negundo** | exotic | perennial | phan | AC | 0 | 1 | 0 | 0 | 0.01 | 0 |
| **Achillea asplenifolia** | native | perennial | herb | DT | 1 | 0 | 3 | 1 | 0 | 2.2 |
| **Achillea collina** | native | perennial | herb | DT | 1 | 2 | 0 | 0.01 | 1.6 | 0 |
| **Agrimonia eupatoria** | native | perennial | herb | DT | 1 | 2 | 0 | 1 | 1.6 | 0 |
| **Agrostis stolonifera** | native | perennial | graminoid | C | 0 | 0 | 3 | 0 | 0 | 8 |
| **Allium scorodoprasum** | native | perennial | herb | DT | 2 | 1 | 1 | 0.2 | 0.01 | 0.01 |
| **Alopecurus pratensis** | native | perennial | graminoid | C | 4 | 2 | 4 | 23 | 6 | 32 |
| **Althaea officinalis** | native | perennial | herb | DT | 0 | 0 | 1 | 0 | 0 | 0.1 |
| **Anacamptis morio** | native | perennial | herb | G | 0 | 0 | 2 | 0 | 0 | 0.21 |
| **Anacamptis palustris** | native | perennial | herb | G | 0 | 1 | 1 | 0 | 0.2 | 0.01 |
| **Angelica sylvestris** | native | perennial | herb | G | 0 | 0 | 5 | 0 | 0 | 4.61 |
| **Anthriscus sylvestris** | native | perennial | herb | DT | 1 | 0 | 0 | 0.5 | 0 | 0 |
| **Arrhenatherum elatius** | native | perennial | graminoid | DT | 5 | 2 | 0 | 15 | 5 | 0 |
| **Briza media** | native | perennial | graminoid | G | 0 | 0 | 1 | 0 | 0 | 0.01 |
| **Bromus commutatus** | native | annual | graminoid | DT | 1 | 1 | 0 | 18 | 1 | 0 |
| **Bromus sterilis** | native | annual | graminoid | RC | 3 | 1 | 0 | 0.12 | 3 | 0 |
| **Calamagrostis epigeios** | native | perennial | graminoid | RC | 3 | 4 | 2 | 3.01 | 7.1 | 1.01 |
| **Caltha palustris** | native | perennial | herb | G | 0 | 0 | 2 | 0 | 0 | 7 |
| **Calystegia sepium** | native | perennial | herb | DT | 1 | 0 | 6 | 2 | 0 | 8.01 |
| **Carex acutiformis** | native | perennial | graminoid | C | 0 | 1 | 0 | 0 | 3 | 0 |
| **Carex distans** | native | perennial | graminoid | C | 0 | 0 | 3 | 0 | 0 | 20 |
| **Carex flacca** | native | perennial | graminoid | G | 3 | 0 | 1 | 90 | 0 | 5 |
| **Carex hirta** | native | perennial | graminoid | DT | 1 | 3 | 2 | 1 | 0.5 | 3 |
| **Carex panicea** | native | perennial | graminoid | G | 0 | 3 | 3 | 0 | 26 | 44 |
| **Carex spicata** | native | perennial | graminoid | DT | 5 | 3 | 2 | 7.01 | 12 | 40 |
| **Carex tomentosa** | native | perennial | graminoid | G | 4 | 4 | 5 | 61 | 81 | 33.1 |
| **Celtis occidentalis** | exotic | perennial | phan | AC | 2 | 1 | 0 | 0.21 | 0.2 | 0 |
| **Centaurea jacea** | native | perennial | herb | G | 0 | 0 | 0 | 0 | 0 | 0 |
| **Cirsium arvense** | native | perennial | herb | RC | 4 | 0 | 0 | 3.4 | 0 | 0 |
| **Cirsium canum** | native | perennial | herb | G | 1 | 1 | 8 | 1 | 1 | 315 |
| **Colchicum autumnale** | native | perennial | herb | G | 1 | 1 | 2 | 0.2 | 2 | 5.5 |
| **Convolvulus arvensis** | native | perennial | herb | RC | 0 | 0 | 0 | 0 | 0 | 0 |
| **Crataegus monogyna** | native | perennial | phan | G | 5 | 3 | 0 | 8.11 | 2 | 0 |
| **Cruciata pedemontana** | native | annual | herb | G | 0 | 1 | 0 | 0 | 8 | 0 |
| **Dactylis glomerata** | native | perennial | graminoid | DT | 4 | 0 | 1 | 1.12 | 0 | 0.01 |
| **Daucus carota** | native | annual | herb | DT | 4 | 2 | 0 | 1.52 | 1.7 | 0 |

| | | | | | | | | | | |
|---|---|---|---|---|---|---|---|---|---|---|
| **Deschampsia caespitosa** | native | perennial | graminoid | C | 4 | 4 | 6 | 19.5 | 21 | 87 |
| **Dipsacus laciniatus** | native | biennial | herb | W | 3 | 1 | 0 | 9 | 3 | 0 |
| **Eleocharis palustris** | native | perennial | graminoid | C | 0 | 0 | 2 | 0 | 0 | 5 |
| **Elymus repens** | native | perennial | graminoid | RC | 5 | 3 | 0 | 9.51 | 10.01 | 0 |
| **Equisetum sp.** | native | perennial | herb | NA | 0 | 2 | 2 | 0 | 1.1 | 0.02 |
| **Erigeron annuus** | exotic | annual | herb | AC | 0 | 0 | 0 | 0 | 0 | 0 |
| **Erysimum repandum** | native | annual | herb | W | 1 | 0 | 0 | 0.01 | 0 | 0 |
| **Festuca pratensis** | native | perennial | graminoid | C | 0 | 0 | 7 | 0 | 0 | 33.01 |
| **Filipendula vulgaris** | native | perennial | herb | G | 1 | 0 | 0 | 0.1 | 0 | 0 |
| **Galium aparine** | native | annual | herb | W | 2 | 4 | 0 | 0.02 | 2.2 | 0 |
| **Galium mollugo** | native | perennial | herb | G | 2 | 2 | 1 | 0.4 | 1.5 | 0.1 |
| **Galium verum** | native | perennial | herb | DT | 3 | 2 | 1 | 7 | 0.02 | 3 |
| **Geum urbanum** | native | perennial | herb | DT | 1 | 1 | 0 | 0.1 | 0.1 | 0 |
| **Glechoma hederacea** | native | perennial | herb | DT | 0 | 0 | 0 | 0 | 0 | 0 |
| **Hieracium sp.** | native | perennial | herb | NA | 1 | 0 | 0 | 0.2 | 0 | 0 |
| **Holcus lanatus** | native | perennial | graminoid | G | 0 | 1 | 0 | 0 | 1 | 0 |
| **Humulus lupulus** | native | perennial | herb | DT | 0 | 1 | 0 | 0 | 2 | 0 |
| **Inula britannica** | native | biennial | herb | DT | 2 | 3 | 0 | 1.1 | 4.3 | 0 |
| **Iris spuria** | native | perennial | herb | S | 1 | 0 | 0 | 3 | 0 | 0 |
| **Juncus gerardii** | native | perennial | graminoid | C | 0 | 0 | 2 | 0 | 0 | 4 |
| **Lathyrus pratensis** | native | perennial | legume | DT | 0 | 0 | 5 | 0 | 0 | 22 |
| **Lathyrus tuberosus** | native | perennial | legume | W | 4 | 3 | 0 | 33.01 | 26.2 | 0 |
| **Leontodon hispidus** | native | perennial | herb | DT | 3 | 0 | 0 | 3.01 | 0 | 0 |
| **Lotus corniculatus** | native | perennial | legume | DT | 0 | 0 | 2 | 0 | 0 | 0.11 |
| **Lychnis flos-cuculi** | native | perennial | herb | G | 1 | 1 | 0 | 3 | 0.1 | 0 |
| **Lycopus exaltatus** | native | perennial | herb | DT | 2 | 1 | 1 | 0.11 | 0.2 | 2 |
| **Lysimachia nummularia** | native | perennial | herb | DT | 1 | 1 | 0 | 4 | 0.3 | 0 |
| **Lysimachia vulgaris** | native | perennial | herb | DT | 1 | 0 | 4 | 0.01 | 0 | 11.5 |
| **Medicago lupulina** | native | annual | legume | DT | 1 | 0 | 0 | 2 | 0 | 0 |
| **Medicago minima** | native | annual | legume | G | 2 | 0 | 0 | 0.2 | 0 | 0 |
| **Melilotus officinalis** | native | annual | legume | W | 0 | 0 | 1 | 0 | 0 | 0.2 |
| **Mentha aquatica** | native | perennial | herb | G | 0 | 0 | 5 | 0 | 0 | 10.02 |
| **Molinia caerulea** | native | perennial | graminoid | C | 0 | 0 | 2 | 0 | 0 | 4 |
| **Myosotis sp.** | native | NA | herb | NA | 0 | 0 | 0 | 0 | 0 | 0 |
| **Pastinaca sativa** | native | perennial | herb | DT | 0 | 0 | 1 | 0 | 0 | 0.1 |
| **Phragmites australis** | native | perennial | graminoid | C | 5 | 5 | 7 | 46 | 22 | 45 |
| **Plantago major** | native | perennial | herb | W | 1 | 0 | 0 | 0.3 | 0 | 0 |
| **Plantago media** | native | perennial | herb | DT | 0 | 0 | 2 | 0 | 0 | 0.3 |
| **Poa pratensis** | native | perennial | graminoid | G | 5 | 1 | 6 | 40 | 2 | 33 |
| **Poa trivialis** | native | perennial | graminoid | DT | 5 | 6 | 1 | 84.1 | 18.21 | 1 |
| **Poaceae** | native | perennial | graminoid | NA | 1 | 0 | 1 | 0.01 | 0 | 1 |
| **Potentilla reptans** | native | perennial | herb | DT | 2 | 0 | 2 | 4 | 0 | 8.01 |
| **Prunus spinosa** | native | perennial | phan | C | 6 | 4 | 0 | 39.12 | 9 | 0 |
| **Pulicaria dysenterica** | native | annual | herb | DT | 1 | 0 | 0 | 1 | 0 | 0 |
| **Pyrus pyraster** | native | perennial | phan | G | 2 | 1 | 0 | 4 | 1 | 0 |
| **Ranunculus acris** | native | perennial | herb | G | 4 | 4 | 3 | 2.1 | 4.01 | 5 |

| | | | | | | | | | | |
|---|---|---|---|---|---|---|---|---|---|---|
| **Ranunculus repens** | native | perennial | herb | DT | 1 | 0 | 4 | 0.01 | 0 | 41 |
| **Rhamnus catharticus** | native | perennial | phan | G | 5 | 4 | 0 | 6.61 | 6 | 0 |
| **Rosa canina** | native | perennial | phan | DT | 1 | 0 | 0 | 0.01 | 0 | 0 |
| **Rubus caesius** | native | perennial | phan | DT | 7 | 6 | 0 | 30.5 | 50 | 0 |
| **Rumex sp.** | native | NA | herb | NA | 1 | 0 | 0 | 0.3 | 0 | 0 |
| **Salix aurita** | native | perennial | phan | S | 0 | 1 | 0 | 0 | 2 | 0 |
| **Scirpus sp.** | native | perennial | graminoid | NA | 0 | 0 | 2 | 0 | 0 | 1.51 |
| **Serratula tinctoria** | native | perennial | herb | G | 1 | 0 | 3 | 0.2 | 0 | 6.2 |
| **Solidago gigantea** | exotic | perennial | herb | AC | 8 | 8 | 0 | 401 | 708 | 0 |
| **Sonchus oleraceus** | native | annual | herb | W | 0 | 0 | 1 | 0 | 0 | 0.2 |
| **Stellaria media** | native | annual | herb | DT | 3 | 0 | 0 | 0.03 | 0 | 0 |
| **Symphytum officinale** | native | perennial | herb | G | 0 | 2 | 1 | 0 | 5.5 | 1 |
| **Taraxacum officinale** | native | perennial | herb | RC | 4 | 1 | 5 | 19 | 1 | 1.8 |
| **Trifolium pratense** | native | perennial | legume | DT | 0 | 0 | 3 | 0 | 0 | 2.01 |
| **Tussilago farfara** | native | perennial | herb | DT | 0 | 1 | 0 | 0 | 1 | 0 |
| **Valeriana officinalis** | native | perennial | herb | G | 2 | 2 | 2 | 0.51 | 0.11 | 2.1 |
| **Valerianella locusta** | native | annual | herb | DT | 4 | 0 | 0 | 0.23 | 0 | 0 |
| **Vicia angustifolia** | native | annual | legume | DT | 3 | 1 | 0 | 2.01 | 0.01 | 0 |
| **Vicia cracca** | native | perennial | legume | DT | 5 | 1 | 5 | 52.5 | 6 | 31.5 |
| **Viola sp.** | native | perennial | herb | G | 2 | 0 | 0 | 0.02 | 0 | 0 |
